# *cis*-eQTL mapping of TB-T2D comorbidity elucidates the involvement of African ancestry in TB susceptibility

**DOI:** 10.1101/2022.10.19.512814

**Authors:** Yolandi Swart, Caitlin Uren, Clare Eckold, Jacqueline M Cliff, Stephanus T Malherbe, Katharina Ronacher, Vinod Kumar, Cisca Wijmenga, Hazel M Dockrell, Reinout van Crevel, Gerhard Walzl, Léanie Kleynhans, Marlo Möller

## Abstract

The validation of genome-wide association signals for tuberculosis (TB) susceptibility and the development of type 2 diabetes (T2D) across diverse populations remain problematic. The ancestry-specific variants (coding and non-coding) that contribute to previously identified differentially expressed genes (DEG) in patients with TB, T2D and comorbid TB-T2D, remain unknown. Identifying ancestry-specific expression quantitative trait loci (eQTLs) can aid in distinguishing the most probable disease-causing variants for population-specific therapeutic interventions. Therefore, this study conducted *cis*-eQTL mapping in TB, T2D and TB-T2D patients to identify variants associated with DEG. Both genotyping (Infinium H3A array with ∼2.3 M markers) and RNA sequencing data of 96 complex multi-way admixed South Africans were used for this purpose. Importantly, both global-and local ancestry adjustment were included in statistical analysis to account for complex admixture. Unique gene-variant pairs were associated with TB-T2D on chromosome 7p22 whilst adjusting for Bantu-speaking African ancestry (*PRKAR1B*:rs4464850; P=7.68e-07) and Khoe-San ancestry (*PRKAR1B:*rs117842122; P=3.66e-07). In addition, *IFITM3* (a biomarker for the development of TB) was associated with three SNPs (rs11025530, rs3808990, and rs10896664) on chromosome 11p15 while adjusting for Khoe-San ancestry. Our results also indicated that the upregulation of the *NLRP6* inflammasome is strongly associated with people with TB-T2D while adjusting for Khoe-San ancestry. Three African-specific eGenes (*NLRP6, IFITM3* and *PRKAR1B*) would have been missed if local ancestry adjustment was not conducted. This study determined a list of ancestry-specific eQTLs in TB-T2D patients that could potentially guide the search for new therapeutic targets for TB-T2D in African populations.

**Author Summary:** The limitation of genome-wide association study (GWAS) is that the particular biological pathway impacted by a variant might not be evident. eQTL mapping can be conducted to determine the impact that a genetic variant might have on the expression of a specific gene in a biological pathway. In this study the use of *cis*-eQTL mapping was explored to elucidate the underlying genetic variants that regulate gene expression between TB-T2D and T2D patients, and between TB patients and healthy controls with multi-way genetic admixture from South Africa. Using RNA sequencing data and newly genotyped dataset of 96 individuals (Illumina Infinium H3Africa array with ∼2.5 M markers), we were able to identify ancestry-specific eQTLs. eQTLs of indigenous Khoe-San ancestral origin were identified in genetic regions previously implicated in TB and T2D in African populations. If local ancestry was not incorporated in the *cis*-eQTL mapping analysis these important African-specific eQTLs would have been missed. Our results provide a list of possible ancestry-specific causal variants associated with TB-T2 comorbidity that could guide the search for new therapeutic targets for African-specific populations. Including populations with complex ancestry and admixture in genetic studies is necessary to improve the quality of genetic research in sub-Saharan African groups.

## Introduction

The dual burden of tuberculosis (TB) and type 2 diabetes (T2D) is a global health problem [1]. Worldwide, an estimated 10 million cases of TB, caused by *Mycobacterium tuberculosis* (*M.tb*), were reported in 2020 [2]. The World Health Organisation (WHO) estimated that African countries accounted for 25% of the estimated 10 million cases of TB, with South Africa at the epicentre of the TB epidemic. More than 15% of all TB patients are estimated to have diabetes which equates to approximately 1.5 million people who require directed therapy and follow-up treatments to manage both diseases [3]. Currently, there is a lack of multidisciplinary approaches to develop therapeutic interventions for infectious and non-communicable diseases in Africa.

Over the past decade, the diabetes prevalence has increased in low- and middle-income countries, where the TB epidemic is also gaining pace at an alarming rate [3–6]. Almost 80% of individuals with T2D in sub-Saharan Africa are undiagnosed and may pose a substantial threat to TB control efforts [7]. According to the International Diabetes Federation (IDF), the diabetes prevalence in Africa is expected to increase by 48% (28 million people) in 2030 and by 129% (55 million people) in 2045, the highest predicted increase of all the IDF Regions [3]. Furthermore, the corona virus disease-19 (COVID-19) pandemic has adversely affected the global efforts to control both TB and T2D, most notably in low-and middle-income countries with populations of diverse ancestry and admixture.

The co-epidemic of TB and T2D is not confined to South Africa or the African continent. South India (54%), some Pacific Islands (40%), South Korea (26.5%), Texas-Mexico (25%), and Ethiopia (15.8%) also suffer from larger numbers of diabetes-associated TB [3]. Moreover, clinical characteristics of TB-T2D vary considerably between countries, for example, the median glycated haemoglobin (HbA1c) among TB-T2D patients in Indonesia is 11.3%, in Peru 10.6%, in South Africa 10.1% and in Romania 7.4% [4].There are thus clear epidemiological and population-specific genetic disease risk factors contributing to TB-T2D comorbidity.

Distinct differentially expressed gene (DEG) profiles were identified in blood to determine the underlying immunological mechanisms that contribute to TB-T2D comorbidity [8]. RNA sequencing of whole blood identified a reduced type 1 interferon response in both TB-T2D patients and TB patients with intermediate hyperglycaemia compared to TB-only patients. Nonetheless, the focus of the study was to identify biomarkers based on DEG between TB-T2D compared to TB, T2D and healthy controls for diagnostic purposes. Thus, the contribution of ancestry-specific genetic variants (coding or non-coding) to the DEG in TB-T2D patients compared to TB, T2D, and healthy controls remains unknown.

A multi-omics approach, such as Expression Quantitative Trait Loci (eQTL) mapping, can provide important information regarding the underlying biological mechanisms of genetic variants (coding or non-coding variants) by linking these to DEG [9]. eQTLs are genetic variants that are associated with gene expression, either located within a short distance (1 Mega base pairs) on either side of a gene’s transcription starting site (TSS) (*cis*-eQTLs) or located at longer distances (5 Mega base pairs) (trans-eQTLs) [10]. This enables the identification of interindividual regulatory candidate variants of transcription and improves our understanding of the effects of genetic polymorphisms on tissue-specific variability in physiological processes [11]. Consequently, eQTL data can be used to model regulatory networks and provide a better understanding of the underlying phenotypic variation.

The major goal of identifying eQTLs is to reduce the number of candidate causal variants for follow-up verification by functional assays [9]. Once identified, eQTLs can provide invaluable genomic information to enhance the power of future GWAS and assist in identifying the most probable disease-causing variants associated with TB-T2D [12]. Currently, the lack of eQTL mapping studies in populations with southern African ancestry hinders the progress of comparative analysis between South African and other populations in terms of differences in genetic architecture underlying gene expression variation [10].

Given the complex nature of both TB and T2D (onset of disease, progression, and treatment variability), *cis*-eQTL mapping was done on samples from South African patients to identify the most probable candidate population-specific causal variants in TB-T2D compared to T2D-only, and in TB-only patients compared to healthy individuals. This was done to understand the genetic risk factors contributing to TB development in T2D patients and healthy controls. Two ancestry adjustment methods, namely Global Ancestry Adjustment (GlobalAA) and Local Ancestry Adjustment (LocalAA), were used. The unique genetic diversity and admixture present in populations in South Africa facilitated the study of ancestry-specific eQTLs. This involves five ancestries from various continents, with differential exposure to *M.tb* throughout history, contributing to the genomic architecture in the country [13].

## Results

### Population structure and ancestry inference of study population

The summary statistics and distributions of the age, sex, body mass index (BMI), and HbA1c for each ancestry are summarized in the supplementary materials (Table S3, Fig S3-5). As expected, there was a significant difference in HbA1c levels between T2D patients and TB-T2D patients compared to no T2D (*P* value = 4.17e-06).

Cross validation was conducted to identify the correct number of contributing ancestral populations (K=3-8) of the admixed population, before inferring global and local ancestry. The estimations indicated that K=5 had the lowest cross validation error (k=0.419, Table S4) and thus represented the most likely number of contributing ancestral populations in the cohort. Fig 1 represents the global ancestry proportions of all 96 admixed individuals included in the statistical analysis. For more refined global ancestry proportions, RFMix was used to infer local and in turn global ancestry. Bantu-speaking African ancestry contributed ∼40.7% of the average global ancestry, indigenous Khoe-San ∼30.8%, European ancestry ∼19.8%, Southeast Asian ancestry ∼6.9% and East Asian ancestry ∼1.9% (Fig 1). In addition to estimating global ancestry using RFMix, local ancestry estimation was conducted, which involves the inference of ancestry at each genomic locus. The local ancestry represented in karyograms indicates the ancestry of each genomic region from chromosome 1 to 22 (Fig S8). Noticeably, the local ancestry patterns appear to be highly heterogeneous, which is in line with previous studies [13,14]. The successful inference of both global and local ancestry allowed the efficient inclusion of this covariate in the subsequent statistical models.

**Fig 1.**
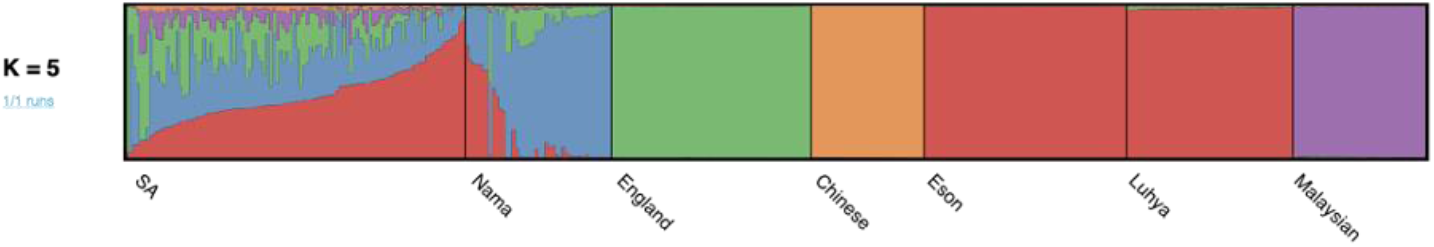
RFMix analysis results using K=5 clusters to infer global ancestry proportions for all 96 admixed South African individuals (SA). The average proportion of Southeast Asian (Malaysian in purple), African (Luhya and Eson in red), East Asian (Chinese in orange), European (England in green), and Khoe-San (Nama in blue) genetic ancestry were 6.9%, 40.7%, 1.9%, 19.8%, and 30.8%, respectively. Displayed populations from left to right on the x-axis: admixed South African individuals from this study (n=96), Khoe-San ancestry (Nama gathered from the European Genome-Phenome archive), Northern and Western European ancestry (GBR from the 1000GP phase 3), East Asian ancestry (CHB from the 1000GP phase 3), Western African ancestry (LWK and MSL from the 1000GP phase 3), and Southeast Asian ancestry (Malaysian from Wong et al.’s 2013 study).

### DEGs amongst TB-T2D-, T2D- and TB patients and healthy controls

In total, 1,581 DEGs were identified when comparing TB-T2D and T2D patients. 178 DEGs were identified between TB patients and healthy controls (Table S5). When quantifying the number of DEGs it became apparent that individuals with preT2D (no TB) had no distinct phenotype compared to T2D patients and healthy controls (Tables S5). For this reason, as well as the low sample number, individuals with preT2D were excluded from the eQTL analysis. Since we were interested in investigating the genetic risk factors (identified through DEG) contributing to TB development in T2D patients and healthy controls, TB-IH patients were excluded from the analysis. Furthermore, DEG analysis of TB-IH patients compared to healthy controls, preT2D and T2D only, were previously reported (Table S5) [8].

Although different DEGs analysis methods (edgeR, limma and voom in R versus DESeq2) were used, this study validated the results on the same cohort as presented in Eckold *et al*. More specifically, overlapping DEGs (across the two studies) include, *BATF2, SOC3, Septin 4, ANKRD22, C1QA, B, C* and *GBP5* when comparing TB patients and healthy controls (Table S6 and S7).

### Ancestry-specific eQTLs identified between TB patients and healthy controls

The eGenes (of which gene expression is associated with at least one genetic variant) and the eQTLs with the corresponding assigned ancestry for each statistical analysis are shown in Table 1. In total, five significant eGenes *(P* value*<*1e-06) were identified. Two of these were identified using LocalAA and three with GlobalAA. Notably, one eGene of Khoe-San origin (ENSG00000269981.1) was identified using GlobalAA. This eGene is located on chromosome 1 and has one transcript which is a splice variant, with no known biological function. Four eQTLs were associated with this eGene (rs2088212; rs2088210; rs10916169; rs903697) and appear to be in linkage with each other (Table 1).

**Table 1.**
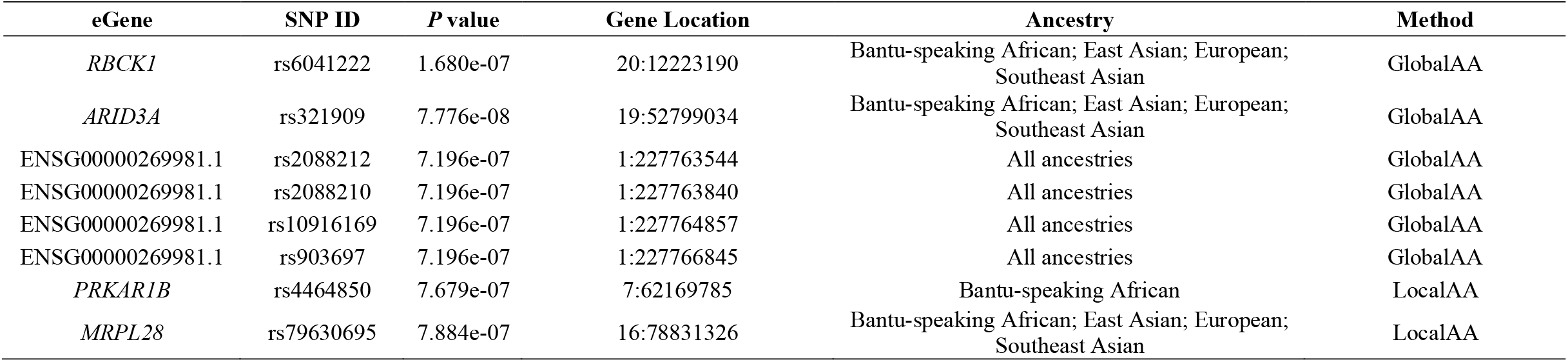
Unique eGenes significantly associated (*P* value <1e-06) with TB patients compared to healthy controls for each ancestry using GlobalAA and LocalAA.

| eGene | SNP ID | $P$ value | Gene Location | Ancestry | Method |
| --- | --- | --- | --- | --- | --- |
| *RBCK1* | rs6041222 | 1.680e-07 | 20:12223190 | Bantu-speaking African; East Asian; European; Southeast Asian | GlobalAA |
| *ARID3A* | rs321909 | 7.776e-08 | 19:52799034 | Bantu-speaking African; East Asian; European; Southeast Asian | GlobalAA |
| ENSG00000269981.1 | rs2088212 | 7.196e-07 | 1:227763544 | All ancestries | GlobalAA |
| ENSG00000269981.1 | rs2088210 | 7.196e-07 | 1:227763840 | All ancestries | GlobalAA |
| ENSG00000269981.1 | rs10916169 | 7.196e-07 | 1:227764857 | All ancestries | GlobalAA |
| ENSG00000269981.1 | rs903697 | 7.196e-07 | 1:227766845 | All ancestries | GlobalAA |
| *PRKAR1B* | rs4464850 | 7.679e-07 | 7:62169785 | Bantu-speaking African | LocalAA |
| *MRPL28* | rs79630695 | 7.884e-07 | 16:78831326 | Bantu-speaking African; East Asian; European; Southeast Asian | LocalAA |

Using LocalAA, the eGene protein Kinase cAMP-Dependent Type I Regulatory Subunit Beta (*PRKAR1B*) was identified. An eQTL (rs4464850) was identified to affect the expression of *PRKAR1B* when comparing TB patients and healthy controls whilst adjusting for Bantu-speaking African ancestry. Interestingly, an eQTL (rs117842122) affecting the expression of *PRKAR1B* was also identified whilst adjusting for Khoe-San ancestry, when comparing TB-T2D to T2D patients. This suggests that this eGene may be implicated in TB progression in T2D patients and healthy controls whilst adjusting for Bantu-speaking African ancestry and Khoe-San ancestry.

Although there are examples where one genetic variant affects the expression of a gene, previous studies suggest that it is more likely that multiple variants affect the expression of a gene [15]. In support of this, our data shows that two eQTLs (rs321909 and rs12459238) both appear to affect the expression of *ARID3A* (AT-Rich Interaction Domain 3A) while adjusting for East Asian ancestry (Table 2). In addition, *ARID3A* was also identified when comparing TB-T2D to T2D patients while adjusting for Khoe-San ancestry. This suggests that this eGene may be implicated in TB progression in both T2D patients and healthy controls. All gene-variant pairs identified with a *P* value threshold of < 1e-04 are summarized in table S8 for TB patients compared to healthy controls.

**Table 2.** Differential lead SNPs (*P* value <1e-06) for the same eGene for TB patients compared to healthy controls using GlobalAA and LocalAA.

| eGene ID | SNP ID | $P$ value | Gene Location | Ancestral Origin | Method |
| --- | --- | --- | --- | --- | --- |
| *SLC6A12* | rs17009851 | 2.341e-07 | 12:83084188 | Bantu-speaking African; European ; East Asian ; Southeast Asian | GlobalAA |
| *ARID3A* | rs321909 | 7.776e-08 | 19:52799034 | East Asian | GlobalAA |
| *SLC6A12* | rs17010578 | 1.460e-08 | 12:83457942 | Bantu-speaking African; European ; East Asian ; Southeast Asian | LocalAA |
| *ARID3A* | rs12459238 | 2.447e-07 | 19:16593359 | East Asian | LocalAA |

Common pathways across eGenes were investigated using GO analysis. When only including eGenes with a *P* value <1e-06, no statistically significant (FDR corrected < 0.05) GO results were observed. However, when decreasing the cut-off to a less stringent value of <1e-04, statistically significant results were observed. Similar results were obtained previously by Eckold *et al*., with genes involved in the type 1 interferon (IFN) signalling pathway, cellular response to type 1 IFN and the IFN-alpha/beta signalling pathways between TB patients and healthy controls while adjusting for Khoe-San ancestry (Table S10). In addition, a NOD-like receptor signalling pathway was identified between TB patients and healthy controls while adjusting for Khoe-San ancestry (Table S10).

### Ancestry-specific eQTLs identified between TB-T2D and T2D patients

The eGenes and the eQTLs with the corresponding assigned ancestry for each statistical analysis is shown in Table 3. In total, four significant (*P* value <1e-06) eGenes were identified using LocalAA. Three eGenes were identified while adjusting for of Khoe-San ancestry and one while adjusting for East Asian ancestry. An eQTL (rs346066) of Khoe-San ancestry origin, affecting the expression of a long non-coding RNA (*LINC01002*) was identified while adjusting for Khoe-San ancestry. Two eGenes (*PRKAR1B* and *ARID3A*), were identified between TB-T2D and T2D patients as well as TB patients and healthy controls while adjusting for Khoe-San ancestry, but were associated with different eQTLs (rs117842122 and rs56369375). Importantly, both eGenes would have been missed if LocalAA was not used. An eQTL (rs2571075) affecting the expression of the ATP Binding Cassette Subfamily A Member 7 (*ABCA7)* was identified while adjusting for East Asian ancestry.

**Table 3.** Unique eGenes significantly associated (*P* value <1e-06) with TB-T2D patients compared to T2D patients for each ancestry using GlobalAA and LocalAA.

| Gene | SNP ID | $P$ value | Gene Location | Ancestral Origin | Method |
| --- | --- | --- | --- | --- | --- |
| *LINC01002 LncRNA* | rs346066 | 9.523e-07 | 19:44217836 | Khoe-San | LocalAA |
| *PRKAR1B* | rs117842122 | 3.655e-07 | 7:76842468 | Khoe-San | LocalAA |
| *ARID3A* | rs56369375 | 7.489e-07 | 19:52770105 | Khoe-San | LocalAA |
| *ABCA7* | rs2571075 | 6.980e-07 | 19:44873148 | East Asian | LocalAA |

Multiple eQTLs affecting the expression of Golgi-associated secretory casein pathway kinase (*FAM20C*), were identified (Table 4). An additional eQTL (rs12531478) affecting the expression of *FAM20C* was identified using LocalAA while adjusting for Bantu-speaking African, East Asian and Khoe-San ancestry. Interestingly, Khoe-San ancestry seems to be associated with *FAM20C* when using LocalAA, but not GlobalAA. Two eQTLs (rs35219837 and rs1186214) affecting the expression of Post-Glycosylphosphatidylinositol Attachment to Protein 6 (*PGAP6*), were identified when comparing TB-T2D and T2D patients while adjusting for Khoe-San ancestry. Three eQTLs (rs11025530, rs3808990 and rs10896664) affecting the expression of NOD-like receptor family Pyrin Domain Containing 6 protein (*NLRP6*), were identified when comparing TB-T2D and T2D patients while adjusting for Khoe-San ancestry. Three eQTLs (rs55970487, rs77247842 and rs12282149) affecting the expression of the IFN-induced transmembrane protein 3 (*IFITM*3), were identified when analysing TB-T2D and T2D patients while adjusting for Khoe-San ancestry. Three eQTLs (rs759202, rs113496159 and rs17677328) affecting the expression of mitochondrial ribosomal protein L28 located (*MRPL28*), were also identified when comparing these two patient groups while adjusting for Khoe-San ancestry. All gene-variant pairs identified with a *P* value threshold of <1e-04 are summarized in table S9 for TB-T2D vs T2D patients.

**Table 4.** Differential lead SNPs (*P* value <1e-06) for the same eGene for TB patients compared to healthy controls using GlobalAA and LocalAA.

| Gene ID | SNP ID | $P$ value | Gene Location | Ancestral Origin | Method |
| --- | --- | --- | --- | --- | --- |
| *FAM20C* | rs11763876 | 7.169e-09 | 7:7249747 | Bantu-speaking African ; East Asian | GlobalAA |
| *FAM20C* | rs78423890 | 7.169e-09 | 7:17071336 | Bantu-speaking African ; East Asian | GlobalAA |
| *FAM20C* | rs57549526 | 7.169e-09 | 7:30198960 | Bantu-speaking African ; East Asian | GlobalAA |
| *FAM20C* | rs62460527 | 7.169e-09 | 7:44908078 | Bantu-speaking African ; East Asian | GlobalAA |
| *FAM20C* | rs183319053 | 7.169e-09 | 7:44910466 | Bantu-speaking African ; East Asian | GlobalAA |
| *FAM20C* | rs12616494 | 7.169e-09 | 7:44910682 | Bantu-speaking African ; East Asian | GlobalAA |
| *FAM20C* | rs188203968 | 7.169e-09 | 7:44911023 | Bantu-speaking African ; East Asian | GlobalAA |
| *FAM20C* | rs62460528 | 7.169e-09 | 7:44912066 | Bantu-speaking African ; East Asian | GlobalAA |
| *FAM20C* | rs115428191 | 7.169e-09 | 7:54748201 | Bantu-speaking African ; East Asian | GlobalAA |
| *FAM20C* | rs57549526 | 4.107e-09 | 7:30198960 | Khoe-San | GlobalAA |
| *MRPL28* | rs759202 | 3.817e-08 | 16:5114437 | Khoe-San | GlobalAA |
| *PGAP6* | rs35219837 | 1.355e-07 | 16:84193640 | Khoe-San | GlobalAA |
| *NLRP6* | rs11025530 | 6.415e-07 | 11:20383324 | Khoe-San | GlobalAA |
| *NLRP6* | rs3808990 | 6.415e-07 | 11:20384800 | Khoe-San | GlobalAA |
| *IFITM3* | rs12282149 | 8.072e-07 | 11:88092069 | Khoe-San | GlobalAA |
| *FAM20C* | rs12531478 | 2.266e-09 | 7:15239894 | Bantu-speaking African | LocalAA |
| *FAM20C* | rs12531478 | 1.242e-09 | 7:15239894 | East Asian | LocalAA |
| *FAM20C* | rs12531478 | 1.237e-09 | 7:15239894 | Khoe-San | LocalAA |
| *MRPL28* | rs113496159 | 9.156e-07 | 16:76054857 | Khoe-San | LocalAA |
| *MRPL28* | rs17677328 | 9.156e-07 | 16:76055758 | Khoe-San | LocalAA |
| *PGAP6* | rs11862144 | 8.888e-08 | 16:54507876 | Khoe-San | LocalAA |
| *NLRP6* | rs10896664 | 5.175e-08 | 11:57707387 | Khoe-San | LocalAA |
| *IFITM3* | rs77247842 | 3.150e-07 | 11:83811362 | Khoe-San | LocalAA |
| *IFITM3* | rs55970487 | 3.150e-07 | 11:83816191 | Khoe-San | LocalAA |

Common pathways across eGenes were investigated using GO analysis. When only including eGenes with a *P* value <1e-06, no statistically significant (FDR corrected < 0.05) GO results were observed. However, when decreasing the cut-off to a less stringent value of <1e-04, statistically significant results were observed. GO analysis indicated a possible upregulation of genes in lung tissue and downregulation of genes in adipose tissue while adjusting for Khoe-San ancestry (Fig S9 and S10). Multiple transcription factors were identified while adjusting for Southeast Asian ancestry. This suggests that Southeast Asian ancestry may have a different biological pathway that drives the development of TB in healthy individuals compared to the other four ancestries in this study. Comparable results were observed between TB-T2D and T2D patients, and TB patients and healthy controls, with genes clustering together in the IFN alpha-beta signalling pathway and NOD-like signalling pathway in both comparisons. This could indicate that both pathways contribute to TB development in T2D patients and healthy controls.

Interestingly, some eGenes overlapped in the GO analysis for both phenotypes (TB patients compared to healthy controls and TB-T2D patients compared to T2D) and clustered in the same genetic regions (Fig 2). *FAM20C* and *PRKAR1B* are both located on chromosome 7p22, *ARID3A* and *ABCA7* overlap on chromosome 19p13 and *NLRP6* and *IFITM3* overlap on chromosome 11p15. This may indicate genetic regions of interest in the context of TB-T2D comorbidity.

**Fig 2.**
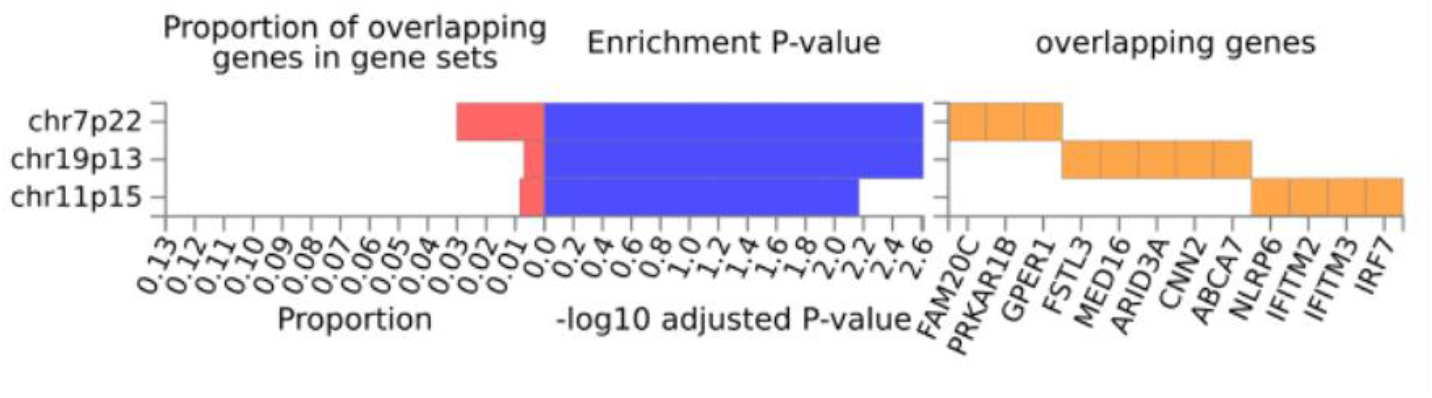
GO analysis of common pathways across significant eGenes for both phenotypes (TB patients compared to healthy controls and TB-T2D patients compared to T2D). The proportion of significant eGenes overlapping between the two phenotypes on similar genetic regions are showcased on the far left, with their corresponding overlapping eGenes on the far-right hand side. The enrichment p-value are indicated in the middle, with chromosome 7p22 and chromosome 19p13 with the strongest enrichment p-value for overlapping eGenes.

## Discussion

Given the absence of TB-T2D comorbidity studies investigating complex multi-way admixed South African populations, this study aimed to identify ancestry-specific eQTLs that contribute to the progression of TB in healthy individuals and T2D patients. Due to the complex multi-way admixed nature of the South African populations, this study used two ancestry adjustment methods (GlobalAA and LocalAA). To our knowledge, this is the first study to link ancestry-specific genetic variants responsible for gene expression in TB, T2D, and TB-T2D patients.

An eQTL (rs4464850) affecting the expression of *PRKAR1B*, was identified when comparing TB patients with healthy controls while adjusting for Bantu-speaking African ancestry. Interestingly, when comparing TB-T2D patients with T2D patients, an eQTL (rs117842122) was identified using LocalAA while adjusting for Khoe-San ancestry. *PRKAR1B* encodes for an important protein kinase regulating the subunit of cyclic AMP-dependent protein kinase A (PKA). *PRKAR1B* is mostly responsible for the cyclic adenosine monophosphate (cAMP)-dependent protein kinase (PKA) signalling pathway which is key in regulating energy balance, glucose homeostasis, and lipid metabolism [16]. A decrease in PKA activity indicated improved lipid profiles in a cohort of obese and overweight African American youths and suggests that an increase in PKA activity may contribute to obesity and insulin resistance [16]. This implies that the upregulation of *PRKAR1B* increases PKA signalling molecules and other proteins regulated by the cAMP signalling pathway and may be associated with T2D and other obesity-related comorbidities in this cohort of African ancestry origin. Notably, only African ancestry was associated with this eGene (*PRKAR1B*) and both eQTLs (rs117842122 and rs4464850) affecting the expression of *PRKAR1B*, would have been missed if only GlobalAA was used in statistical analysis.

A similar trend was observed for the eGene, *ARID3A*. Two eQTLs (rs12459238 and rs321909) affecting the expression of *ARID3A*, were identified when comparing TB patients with healthy controls while adjusting for East Asian ancestry. Similar to *PRKAR1B*, when comparing TB-T2D with T2D patients, an eQTL (rs56369375) of Khoe-San ancestry origin was identified using LocalAA while adjusting for Khoe-San ancestry. *ARID3A* is a potential biomarker for TB diagnosis and treatment response in peripheral blood of TB patients [17]. In addition, *ARID3A* plays an important role in immune responses against intracellular pathogens by controlling cell cycle progression via the RB1/E2F1 pathway and is essential for the development of B-cells. Since both eGenes (*PRKAR1B* and *ARID3A*) were identified in both phenotypes, it suggests that these two eGenes contribute to TB progression in both T2D patients and healthy controls.

eQTLs affecting the expression of *PGAP6, NLRP6* and *IFITM3*, were identified when comparing TB-T2D and T2D patients while adjusting for Khoe-San ancestry. *IFITM3* forms part of a four-gene signature that is able to distinguish active TB patients from healthy controls [18] and is also one of the seventeen TB biomarkers in UK and Indian populations [19]. *IFITM3*, localized on chromosome 11p15, is a genetic region that has previously been linked to TB susceptibility[20,21], and paediatric TB patients of Han Chinese origin [22,23]. *PGAP6* is upregulated in gestational diabetes patients and is inversely correlated with gene expression in type 1 diabetes [24]. *NLRP6* mediates inflammasome activation in response to various pathogen-associated signals, as part of the sensor component of the *NLRP6* inflammasome [25–27].

Inflammasomes play a critical role in innate immunity and inflammation by assembling in the cytosol and acting as a recognition receptor to bind pathogens and other damage-associated signals [26,28]. Interestingly, the dysregulation of inflammasomes has been found to be involved in the pathogenesis of chronic inflammatory diseases such as multiple sclerosis, atherosclerosis, T2D and obesity [29]. When pro-inflammatory macrophages infiltrate the pancreatic islets of T2D patients, it drives the production of IL-1beta via the *NLRP3* inflammasome [30]. Beta-cell proliferation is initially favoured by low concentrations of IL-1beta, however, chronically elevated levels of IL-1beta may lead to beta-cell failure [31]. In contrast, administration of an IL-1 receptor antagonist, improves glucose tolerance, beta-cell function, and systematic inflammation in humans. Moreover, metabolites produced by intestinal microbiota may drive the development of insulin resistance in obesity and T2D by initiating an inflammatory response via *NLRP6* [31]. Henao-Mejia *et al*. confirmed the observation that dysbiosis of the microbiota is linked to metabolic diseases in *NLRP6* mutant mice [32]. Furthermore, *NLRP6* mutant mice have enhanced activation of MAPK and NF-kB signaling via the activation of Toll-like receptors (TLR) and therefore an increased number of immune cells in circulation [33].

*NLRP6* is a negative regulator of inflammatory signaling and NF-kB signaling in response to bacterial pathogens in myeloid cells ^7^. Therefore, *NLRP6* expression may prevent clearance of both gram-positive and gram-negative bacterial pathogens. Inflammasome inhibitors that target the polymorphisms of *NLRP6* in TB-T2D patients may provide new means of therapeutic interventions for patients with Khoe-San ancestry and may help alleviate the dual burden of both diseases.

Interestingly, both *NLRP6* and *IFITM3* are located on chromosome 11p15. This region was associated with multiple facets of innate and adaptive immune responses. The *IRF7* gene is located in this region and was associated with developing severe TB [34]. Chromosome 11 was identified in a meta-analysis and a trans-ethnic fine-mapping study to be associated with TB and includes the involvement of the WT1 signalling pathway [20,35]. Interestingly, the *KCNQ1* gene cluster maps within the 11p15.5 imprinted domain and variants intronic to *KCNQ1* influence diabetes susceptibility which is maternally inherited during early development [36]. Furthermore, *KNCNQ1* has been established as a candidate susceptibility gene for T2D and influences the K 7.1 voltage-gated potassium channel subunit located in human beta cells [37]. This evidence points to the involvement of chromosome 11p15 in the development of T2D in individuals of Khoe-San ancestry origin.

An eQTL (rs2571075) affecting the expression of *ABCA7*, was identified when comparing TB-T2D with T2D patients while adjusting for East Asian ancestry. *ABCA7* plays a role in multiple biological processes such as lipid homeostasis, macrophage-mediated phagocytosis, binds APOA1, apolipoprotein-mediated phospholipid efflux from cells and possibly mediates cholesterol efflux [38–41]. The impact on TB or T2D remains unclear, however, it is involved in the phagocytosis of apoptotic cells by macrophages. Macrophage phagocytosis is stimulated by APOA1 or APOA2 upon the stabilization of *ABCA7* [42].

The eGenes that overlapped in certain genetic regions were previously associated with TB susceptibility. Two eGenes (*FAM20C* and *PKRA1B*), are both located on chromosome 7p22. This region is associated with TB susceptibility in a Ugandan population [43]. Likewise, *ARID3A* and *ABCA7* are located on chromosome 19p13. This region was associated with TB susceptibility and linked with the *CD209* gene. This gene – coding for Dendritic Cell-Specific ICAM3-Grabbing Non-integrin (DC-SIGN), is one of the major receptors for *M.tb* on human dendritic cells. A relatively large number of studies evaluated the association between *CD209* polymorphisms (*-336A/G, -871A/G*) and TB risk, but the results have been inconsistent due to limited sample sizes and different studies populations [44,45].

It has been hypothesized that the inclusion of local ancestry in eQTL mapping increases the power to identify novel ancestry-specific eQTLs [46,47]. Most of the eGenes and the eQTLs with their corresponding assigned ancestry would not have been identified if LocalAA was not used. Furthermore, the eQTL (rs12531478) of Khoe-San ancestry origin, affecting the expression of *FAM20C*, was only elucidated once applying LocalAA. This indicates that important indigenous African-specific genetic variants could be missed when only global ancestry is used to account for population structure in complex admixed South African individuals.

Given our modest sample size, findings should be validated in ethnically similar cohorts. Furthermore, whole-genome sequencing could help identify structural variants (small insertions, deletions (indels), and larger structural variations, such as duplications, inversions, and translocations involved in TB-T2D comorbidity. Additionally, future studies should investigate the possible role of methylation (ATAC sequencing) on the DEG, since multiple mechanisms (not only genetic variants) could influence gene expression. *cis*-eQTLs only identify nearby variants located near DEG (1Mb upstream or downstream). Although the extent of involvement of *trans*-eQTLs is still uncertain [9,48], it would still be worthwhile to investigate. In addition, genes that are located near GWAS-significant hits from previous studies that are also identified to be eGenes may be candidate causal genes. Therefore, the different lead variants identified for each ancestry for the same eGenes should be included in future studies to compare it to previous GWAS hits for TB and T2D. This will assist with the prioritization of GWAS hits for inclusion in follow-up functional studies. Together, gene-variant pairs can give supporting evidence (genetic information) for GWAS hits.

In conclusion, incorporating local ancestry in *cis*-eQTL mapping enabled the identification of ancestry-specific eQTLs between TB-T2D and T2D patients, as well as between TB patients and healthy controls. Furthermore, a list of possible candidate disease-causing variants was identified between TB-T2D and T2D patients, as well as between TB patients and healthy controls which could be functionally validated. This could facilitate the early identification of T2D patients at risk of developing TB and may improve the health of complex multi-way admixed South Africans.

## Material and methods

### Ethics Approval and sample collection

Sample collection (protocol number N13/05/064) and the research presented here (S20/02/041) were both approved by the Health Research Ethics Committee (HREC) of the Faculty of Medicine and Health Sciences, Stellenbosch University. The research was conducted according to the principles expressed in the Declaration of Helsinki (2013). Written informed consent was obtained from all study participants before recruitment and blood collection.

Healthy controls, T2D patients without TB as well as TB patients with and without T2D were recruited between December 2013 and February 2016 from communities located in the Northern Suburbs of Cape Town, South Africa as part of the TANDEM study [4]. TB patients were either bacteriologically confirmed (culture positive) or diagnosed by GeneXpert. All participants were between the age of 18 and 70 years and tested negative for HIV. Participants were excluded from the study if they were already on TB medication, receiving steroids therapy, had other serious conditions including cancer, were pregnant, or using excessive amounts of alcohol or illicit drugs. Gestational or steroid-induced diabetes was also excluded. Participants were classified into different groups based on reference laboratory HbA1c levels. Healthy controls (n=23) and TB patients without T2D (TB only; n=10) had an Hb1Ac <5.7% mmol/L. PreT2D (n=6) and TB patients with intermediate hyperglycaemia (TB-IH; n=19) had an Hb1Ac of 5.7% to < 6.5% mmol/L, T2D patients (n=28), including TB patients with T2D (TB-T2D; n=10) had an Hb1Ac ≥ 6.5 mmol/L.

### Genotype data

DNA was extracted for 96 individuals using the Qiagen Blood Midi kit (Qiagen, Germany) as recommended by the manufacturer. RNA sequencing data of the same 96 study participants were also available [8]. Genotype data was generated for all individuals using the Illumina Infinium Human, Hereditary and Health (H3Africa) Consortium Array v2 (comprising ∼2.3 million markers) at the Centre for Proteomic and Genomic Research, South Africa. The H3Africa array was designed to efficiently capture and characterise the genetic diversity in Africa [49]. GenomeStudio v2.04 (Illumina, Miami, United States) was used to calculate intensity scores and call common variants (MAF ≥ 5%) [50]. The software *zCall* was used to recall variants (MAF > 1% and < 5%) [51]. Variants called by GenomeStudio were exported as PLINK formatted files for downstream data analysis.

### Quality control and imputation of genotype data

Quality control of the raw genotype data was done using a reproducible snakemake pipeline (https://github.com/hennlab/snake-SNP_QC) to filter out low-quality samples and SNPs [52]. Quality control and filtering parameters applied to the raw genotypes are indicated in Fig S1.

GenomeHarmonizer version 3 [53] was used to align the data to the 1000 Genomes Phase 3 reference panel (Human genome build 37) [54], to update SNP IDs and remove any variants, not in the reference panel. A minimum linkage disequilibrium (LD) of 0.3 with at least three flanking variants was required for strand alignment. A secondary minor allele frequency (MAF) alignment was also used at a threshold of 5%. Finally, the minimum posterior probability to call genotypes in the input data was left at the default value of 0.4.

After filtering and quality control of the genotypic data, it was converted from a PLINK file format to Variant Call Format (VCF) using PLINK v2.0 [55]. The Sanger Imputation Server was used for phasing, using SHAPEIT2 [56], followed by imputation using the Positional Burrows-Wheeler Transformation (PBWT) algorithm and the African Genome Resource Panel [57]. VCF files were downloaded from the online server after imputation and converted to PLINK ped/map files using a genotyping threshold of 0.7 (PLINK command: -vcf-min-gp command and -output-missing-genotype N).

The *UCSC liftOver* was used to convert the phased, imputed H3Africa genetic data from reference genome Human genome build 37 (hg19) to Human genome build 38 (hg38) to ensure compatibility with the gene expression data required to conduct the eQTL mapping [58]. After performing the imputation, phasing, quality control and filtering, the final dataset comprised of 4 224 844 variants and 96 individuals (summarized in Table S1 and Fig S2).

### Global Ancestry Inference

The genotype data was merged with the appropriate source populations (summarized in Table S2) using PLINK v2.0 [55], to generate input files required for global and local ancestry inference. After merging, all individuals missing more than 10% of the genotypes were removed, SNPs with more than 3% missing data were excluded and a Hardy-Weinberg Equilibrium (HWE) filter of 0.01 was used. The software KING was used to determine relatedness between individuals up to 2nd degree relatedness [59].

The software ADMIXTURE was used to investigate the population structure of the cohort and to determine the correct number of contributing ancestries [60,61]. Each SNP in LD was defined as *r*^*2*^ > 0.1 within a 50-SNP sliding window (advanced by 10 SNPs at a time) and was removed for the purpose of computational efficiency. A total of 273,175 autosomal markers remained after LD pruning. Global ancestry was inferred in an unsupervised manner for K=3-8, where K represents the number of contributing ancestral populations. After establishing the correct K number of contributing ancestries through cross-validation, the software RFMix was used to infer global ancestry proportions for downstream statistical analysis (see specific parameters below), since ADMIXTURE is not as accurate as haplotype-based analyses [62].

### Local Ancestry Inference

The software RFMix was used to infer local ancestry [63]. Default parameters were used, except for the number of generations since admixture, which was set to 15, consistent with previous studies [13]. A total of 4,230,650 autosomal variants were included. For each individual, consecutive phased alleles with the local ancestry assignment were collapsed into BED files of haplotype blocks. These local ancestry BED files were then used to count the number of African, Khoe-San, European, Southeast Asian, and East Asian alleles at each SNP.

### Gene expression data

Venous blood was collected using PAXgene Blood RNA tubes (PreAnalytiX). Sample collection occurred at TB diagnosis (baseline) before TB treatment commenced. Total RNA was extracted using the PAXgene Blood miRNA kit (Qiagen, Germany) with the semi-automated QIAcube (Qiagen, Germany) [8]. RNA sequencing was conducted using the NextSeq500 High Output kit v2 (Illumina) for 75 cycles. The polyA tail library preparation method was used and single-end read sequencing was conducted (n=103) [8].

Quality control, filtering and trimming of raw reads were conducted with HTStream v1.3.1 (Releases s4hts/HTStream). Raw RNA sequencing reads were mapped to the human reference genome (release GRCh38) using STAR v2.5.3a with default parameters [64]. Gene-level quantification was performed with STAR Aligner using the GENCODE v34 annotation file and a subsequent counting table was generated and used as input for DEG identification. Quantified gene expression (TPM and raw counts) was filtered and normalized using the R-package edgeR, limma and voom packages.

### *Cis*-eQTL mapping with LocalAA and GlobalAA

An approach similar to that of Zhong *et al*. (2018) and Gay *et al*. (2019) was used to incorporate both global and local ancestry whilst conducting *cis*-eQTL mapping in a multi-way admixed South African population. This method allows for the identification of associations between variants and gene expression for each contributing ancestral population [46,65]. Genome-wide *cis-*eQTL mapping was performed on 96 individuals and 4,230,650 autosomal variants. All analyses were performed independently for each of the five contributing ancestries (Bantu-speaking African, Khoe-San, European, Southeast Asian and East Asian). The normalized gene expression files were used to calculate 15 hidden confounders with PEER [66]. Additional sample-level covariates (age, gender and HbA1c) were also included in the association analysis.

The following linear regression model was fitted for each gene-variant pair (gene *g*, variant *v*):

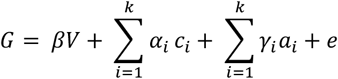

***G*** represents the differential gene expression of gene *g* across all 96 admixed individuals.

***V*** represents the additive effect of alleles at variant *v* (coded as 0,1 or 2).

*β* represents the effect size of the alleles of variant *v* on gene *g* expression.

*α*_*i*_ represents the biological or technical covariate *C*_*i*_ on gene ***g*** expression. This includes age, gender, HbA1c and the PEER hidden confounding factors.

*γ*_*i*_ represents the effect of the ancestry covariate *α*_*i*_ on gene ***g*** expression.

***e*** represents the residual.

Two iterations of this regression were performed for each gene-variant pair.

1. **Global Ancestry Adjustment (GlobalAA):** Adjusting for global ancestry proportions *a*_*i*_ represents the global ancestry proportions of each admixed individual.
2. **Local Ancestry Adjustment (LocalAA):** Adjusting for local ancestry, in which the number of alleles at variant *v* were assigned to a specific ancestry of interest (1 = ancestry of interest; 0 = other ancestries)

If any of the 4,230,650 filtered variants were located within one mega base of the transcription start site, they were included in the association analysis with the gene expression. The *lm()* function in R was used for all regressions performed. An additive genetic effect on gene expression was assumed. The significance of an association was taken to be the two-sided *P* value corresponding to the *t*-statistic of the *β* coefficient estimate. Additionally, the most significant lead eQTLs were identified for each gene, independently for each ancestry adjustment method. To approximate a 5% False Discovery Rate (FDR), a nominal *P* value of 1e-6 to identify significant associations was applied [46]. To discern which biological functions are shared amongst the significant eGenes, gene ontology (GO) and Kyoto Encyclopedia of Genes and Genomes (KEGG) pathway enrichment analyses were done for each ancestry separately. The web-based software g:Profiler was used for this purpose and the default option g:SCS method in g:Profiler was used for multiple testing corrections. Pathways with an adjusted *P* value < 0.05 were reported [67]. Fig S2 summarizes the analysis pipeline used for *cis*-eQTL mapping.

## Acknowledgments

We acknowledge the Centre for High Performance Computing (CHPC), South Africa, for providing computational resources to this research project. The authors thank the clinical staff and the patients at the 4 study sites, as well as Bahram Sanjabi, Desiree Brandenburg-Weening, and Pieter van der Vlies for assistance with the RNA-seq.

## Members of the TANDEM (Concurrent Tuberculosis and Diabetes Mellitus;unravellingg the causal link, and improving care) Consortium

The TANDEM partners and collaborators include: H. Dockrell, J. Cliff, C. Eckold, D. Moore, U. Griffiths, and Y. Laurence (London School of Hygiene and Tropical Medicine, London, United Kingdom); R. Aarnouste, M. Netea; R. van Crevel, C. Ruesen, and E. Lachmandas (Radboud University Medical Center, Nijmegen, The Netherlands); S. Kaufmann, M. Beigier, and R. Golinski (Max Planck Institute for Infection Biology, Berlin, Germany); S. Joosten, T. Ottenhoff, F. Vrieling, and M. Haks (Leiden University Medical Center, Leiden, The Netherlands); G. Walzl, K. Ronacher, S. Malherbe, L. Kleynhans, B. Smith, K. Stanley, G. van der Spuy, A. Loxton, N. Chegou, M. Bosman, L. Thiart, C. Wagman, H. Tshivhula, M. Selamolela, N. Prins, W. du Plessis, I. van Rensburg, and L. du Toit (Stellenbosch University, Cape Town, South Africa); J. Critchley, S. Kerry-Barnard, F. Pearson, and D. Grint (St George’s, University of London, London, United Kingdom); S. McAllister, P. Hill, and A. Verrall (University of Otago, Dunedin, New Zealand); M. Ioana, A. Riza, R. Cioboata, M. Dudau, F. Nitu, I. Bazavan, M. Olteanu, C. Editoiu, A. Florescu, M. Mota, S. G. Popa, A. Firanescu, A. Popa, I. Gheonea, S. Bicuti, A. Lepadat, I. Vladu, D. Clenciu, M. Bicu, C. Streba, A. Demetrian, M. Ciurea, A. Cimpoeru, A. Ciocoiu, S. Dorobantu, R. Plesea, E. L. Popescu, M. Cucu, I. Streata, F. Burada, S. Serban-Sosoi, N. Panduru, and E. Nicoli (University of Craiova, Craiova, Romania); M. Ciontea, I. Capitanescu, M. Olaru, T. Tataru, M. Papurica, I. Valutanu, V. Dubreu, and L. Stamatoiu (Pneumoftisiology Hospital, Gorj, Romania); V. Kumar and C. Wijmenga (University of Groningen, Groningen, The Netherlands); C. Ugarte-Gil, J. Coronel, S. Lopez, R. Limascca, K. Villaizan, B. Castro, J. Flores, and W. Solano (Universidad Peruana Cayetano, Lima, Peru); B. Alisjahbana, R. Ruslami, N. Soetedjo, P. Santoso, L. Chaidir, R. Koesoemadinata, N. Susilawati, J. Annisa, R. Livia, V. Yunivita, A. Soeroto, H. Permana, S. Imaculata, Y. Gunawan, N. Dewi, and L. Apriani (Universitas Padjadjaran, Bandung, Indonesia).

## Author contributions

YS conceived and designed and conducted analysis and wrote the paper. LK provided DNA samples for genotyping and critically assessed manuscript and analysis. CU and MM critically assessed the analysis and manuscript. CE, JMC, STM KR, VK, CW, HMD, RvC, and GW established sample bank and reviewed manuscript.

## Supporting Information

**S1 Fig**. Flow diagram of quality control and filtering parameters applied to raw genotypic data.

**S2 Fig**. Flow diagram of the data analysis pipeline used for *cis-*eQTL mapping.

**S3 Fig**. Distribution of age for all phenotypes.

**S4 Fig**. Distribution of BMI for all phenotypes.

**S5 Fig**. Distribution of HbA1c between all phenotypes.

**S6 Fig**. Ancestry proportions between TB-T2D and T2D patients.

**S7 Fig**. Ancestry proportions between TB patients and healthy controls.

**S8 Fig**. Karyograms representing four individuals’ local ancestry at each genomic region from chromosome 1 to 22.

**S9 Fig. Gene Ontology analysis**. Gene ontology analysis revealed general tissue types in which DEG would most likely occur while adjusting for Khoe-San ancestry only in TB-T2D comorbid patients compared to T2D.

**S10 Fig. Gene ontology analysis**. Gene ontology analysis revealed general tissue types where DEG were mostly up-or-down regulated in TB-T2D comorbid patients compared to T2D in Khoe-San individuals using GlobalAA.

**S1 Table**. Quality control filtering parameters and the total number of variants and/or individuals removed by the filtering command.

**S2 Table**. Ancestral populations included in analysis for ancestry inference.

**S3 Table**. Summary statistics of age, gender, HbA1c, BMI and ancestry proportions.

**S4 Table**. Cross validation error values for K=3-8 ancestral populations

**S5 Table**. Total number of DEG identified for each comparison. Highlighted in green is the two comparisons used for *cis*-eQTL mapping.

**S6 Table**. Top DEG between TB patients and healthy controls. Highlighted genes are the same DEG as previously identified in Eckold *et al*.

**S8 Table**. Differential lead SNPs (<1e-04) for the same eGene for TB-T2D patients compared to T2D patients for all ancestries.

**S9 Table**. Differential lead SNPs (<1e-04) for the same eGene for TB patients compared to Healthy controls for all ancestries.

**S10 Table**. GO between TB patients and Healthy controls of Khoe-San ancestry origin.

**S11 Table**. GO between TB patients and Healthy controls of Southeast Asian ancestry origin.

## References

1. Ngo MD, Bartlett S, Ronacher K. Diabetes-Associated Susceptibility to Tuberculosis: Contribution of Hyperglycemia vs. Dyslipidemia. Microorganisms. 2021;9: 2282. doi:10.3390/microorganisms9112282

2. Global Tuberculosis Report 2021. [cited 22 Nov 2021]. Available: https://www.who.int/teams/global-tuberculosis-programme/tb-reports/global-tuberculosis-report-2021

3. Home, Resources, diabetes L with, Acknowledgement, FAQs, Contact, et al. IDF Diabetes Atlas | Tenth Edition. [cited 19 Jan 2022]. Available: https://diabetesatlas.org/

4. Ugarte-Gil C, Alisjahbana B, Ronacher K, Riza AL, Koesoemadinata RC, Malherbe ST, et al. Diabetes Mellitus Among Pulmonary Tuberculosis Patients From 4 Tuberculosis-endemic Countries: The TANDEM Study. Clin Infect Dis. 2019 [cited 18 Sep 2019]. doi:10.1093/cid/ciz284

5. Tenaye L, Mengiste B, Baraki N, Mulu E. Diabetes Mellitus among Adult Tuberculosis Patients Attending Tuberculosis Clinics in Eastern Ethiopia. In: BioMed Research International [Internet]. 2019 [cited 15 Jan 2020]. doi:10.1155/2019/7640836

6. Lutfiana NC, van Boven JFM, Masoom Zubair MA, Pena MJ, Alffenaar JC. Diabetes mellitus comorbidity in patients enrolled in tuberculosis drug efficacy trials around the world: A systematic review. Br J Clin Pharmacol. 2019;85: 1407–1417. doi:10.1111/bcp.13935

7. Pheiffer C, Pillay-van Wyk V, Joubert JD, Levitt N, Nglazi MD, Bradshaw D. The prevalence of type 2 diabetes in South Africa: a systematic review protocol. BMJ Open. 2018;8: e021029. doi:10.1136/bmjopen-2017-021029

8. Eckold C, Kumar V, Weiner J, Alisjahbana B, Riza A-L, Ronacher K, et al. Impact of Intermediate Hyperglycemia and Diabetes on Immune Dysfunction in Tuberculosis. Clinical Infectious Diseases. 2021;72: 69–78. doi:10.1093/cid/ciaa751

9. Westra H-J, Franke L. From genome to function by studying eQTLs. Biochim Biophys Acta. 2014;1842: 1896–1902. doi:10.1016/j.bbadis.2014.04.024

10. Shang L, Smith JA, Zhao W, Kho M, Turner ST, Mosley TH, et al. Genetic Architecture of Gene Expression in European and African Americans: An eQTL Mapping Study in GENOA. The American Journal of Human Genetics. 2020;106: 496–512. doi:10.1016/j.ajhg.2020.03.002

11. Gay NR, Gloudemans M, Antonio ML, Balliu B, Park Y, Martin AR, et al. Impact of admixture and ancestry on eQTL analysis and GWAS colocalization in GTEx. bioRxiv. 2019; 836825. doi:10.1101/836825

12. Hu Y-J, Sun W, Tzeng J-Y, Perou CM. Proper Use of Allele-Specific Expression Improves Statistical Power for cis-eQTL Mapping with RNA-Seq Data. J Am Stat Assoc. 2015;110: 962–974. doi:10.1080/01621459.2015.1038449

13. Uren C, Kim M, Martin AR, Bobo D, Gignoux CR, van Helden PD, et al. Fine-Scale Human Population Structure in Southern Africa Reflects Ecogeographic Boundaries. Genetics. 2016;204: 303–314. doi:10.1534/genetics.116.187369

14. Swart Y, Uren C, van Helden PD, Hoal EG, Möller M. Local Ancestry Adjusted Allelic Association Analysis Robustly Captures Tuberculosis Susceptibility Loci. Frontiers in Genetics. 2021;12: 2043. doi:10.3389/fgene.2021.716558

15. Spielman RS, Bastone LA, Burdick JT, Morley M, Ewens WJ, Cheung VG. Common genetic variants account for differences in gene expression among ethnic groups. Nat Genet. 2007;39: 226–231. doi:10.1038/ng1955

16. Bloyd M, Settas N, Faucz FR, Sinaii N, Bathon K, Iben J, et al. The PRKAR1B p.R115K Variant is Associated with Lipoprotein Profile in African American Youth with Metabolic Challenges. J Endocr Soc. 2021;5: bvab071. doi:10.1210/jendso/bvab071

17. Xie L, Chao X, Teng T, Li Q, Xie J. Identification of Potential Biomarkers and Related Transcription Factors in Peripheral Blood of Tuberculosis Patients. Int J Environ Res Public Health. 2020;17: 6993. doi:10.3390/ijerph17196993

18. Suliman S, Thompson E, Sutherland J, Weiner Rd J, Ota MOC, Shankar S, et al. Four-gene Pan-African Blood Signature Predicts Progression to Tuberculosis. Am J Respir Crit Care Med. 2018. doi:10.1164/rccm.201711-2340OC

19. Perumal P, Abdullatif MB, Garlant HN, Honeyborne I, Lipman M, McHugh TD, et al. Validation of Differentially Expressed Immune Biomarkers in Latent and Active Tuberculosis by Real-Time PCR. Front Immunol. 2020;11: 612564. doi:10.3389/fimmu.2020.612564

20. Chimusa ER, Zaitlen N, Daya M, Möller M, van Helden PD, Mulder NJ, et al. Genome-wide association study of ancestry-specific TB risk in the South African Coloured population. Hum Mol Genet. 2014;23: 796–809. doi:10.1093/hmg/ddt462

21. Thye T, Owusu-Dabo E, Vannberg FO, van Crevel R, Curtis J, Sahiratmadja E, et al. Common variants at 11p13 are associated with susceptibility to tuberculosis. Nat Genet. 2012;44: 257–259. doi:10.1038/ng.1080

22. Chen C, Zhao Q, Shao Y, Li Y, Song H, Li G, et al. A Common Variant of ASAP1 Is Associated with Tuberculosis Susceptibility in the Han Chinese Population. In: Disease Markers [Internet]. 2019 [cited 3 May 2019]. doi:10.1155/2019/7945429

23. Shen C, Wu X, Jiao W, Sun L, Feng W, Xiao J, et al. A functional promoter polymorphism of IFITM3 is associated with susceptibility to pediatric tuberculosis in Han Chinese population. PLoS One. 2013;8: e67816. doi:10.1371/journal.pone.0067816

24. Starskaia I, Laajala E, Grönroos T, Härkönen T, Junttila S, Kattelus R, et al. Early DNA methylation changes in children developing beta cell autoimmunity at a young age. Diabetologia. 2022;65: 844–860. doi:10.1007/s00125-022-05657-x

25. Hara H, Seregin SS, Yang D, Fukase K, Chamaillard M, Alnemri ES, et al. The NLRP6 Inflammasome Recognizes Lipoteichoic Acid and Regulates Gram-Positive Pathogen Infection. Cell. 2018;175: 1651-1664.e14. doi:10.1016/j.cell.2018.09.047

26. Shen C, Lu A, Xie WJ, Ruan J, Negro R, Egelman EH, et al. Molecular mechanism for NLRP6 inflammasome assembly and activation. Proc Natl Acad Sci U S A. 2019;116: 2052–2057. doi:10.1073/pnas.1817221116

27. Tian XX, Li R, Liu C, Liu F, Yang LJ, Wang SP, et al. NLRP6-caspase 4 inflammasome activation in response to cariogenic bacterial lipoteichoic acid in human dental pulp inflammation. Int Endod J. 2021;54: 916–925. doi:10.1111/iej.13469

28. Ghimire L, Paudel S, Jin L, Jeyaseelan S. The NLRP6 inflammasome in health and disease. Mucosal Immunol. 2020;13: 388–398. doi:10.1038/s41385-020-0256-z

29. Wawrocki S, Druszczynska M. Inflammasomes in Mycobacterium tuberculosis-Driven Immunity. Can J Infect Dis Med Microbiol. 2017;2017: 2309478. doi:10.1155/2017/2309478

30. Zhong Y, Kinio A, Saleh M. Functions of NOD-Like Receptors in Human Diseases. Frontiers in Immunology. 2013;4. Available: https://www.frontiersin.org/article/10.3389/fimmu.2013.00333

31. Scheithauer TPM, Rampanelli E, Nieuwdorp M, Vallance BA, Verchere CB, van Raalte DH, et al. Gut Microbiota as a Trigger for Metabolic Inflammation in Obesity and Type 2 Diabetes. Front Immunol. 2020;11: 571731. doi:10.3389/fimmu.2020.571731

32. Henao-Mejia J, Elinav E, Jin C, Hao L, Mehal WZ, Strowig T, et al. Inflammasome-mediated dysbiosis regulates progression of NAFLD and obesity. Nature. 2012;482: 179–185. doi:10.1038/nature10809

33. Anand PK, Malireddi RKS, Lukens JR, Vogel P, Bertin J, Lamkanfi M, et al. NLRP6 negatively regulates innate immunity and host defence against bacterial pathogens. Nature. 2012;488: 389–393. doi:10.1038/nature11250

34. Cubillos-Angulo JM, Arriaga MB, Melo MGM, Silva EC, Alvarado-Arnez LE, de Almeida AS, et al. Polymorphisms in interferon pathway genes and risk of Mycobacterium tuberculosis infection in contacts of tuberculosis cases in Brazil. International Journal of Infectious Diseases. 2020;92: 21–28. doi:10.1016/j.ijid.2019.12.013

35. Thye T, Owusu-Dabo E, Vannberg FO, van Crevel R, Curtis J, Sahiratmadja E, et al. Common variants at 11p13 are associated with susceptibility to tuberculosis. Nat Genet. 2012;44: 257–259. doi:10.1038/ng.1080

36. Travers ME, Mackay DJG, Dekker Nitert M, Morris AP, Lindgren CM, Berry A, et al. Insights Into the Molecular Mechanism for Type 2 Diabetes Susceptibility at the KCNQ1 Locus From Temporal Changes in Imprinting Status in Human Islets. Diabetes. 2013;62: 987–992. doi:10.2337/db12-0819

37. Asahara S, Etoh H, Inoue H, Teruyama K, Shibutani Y, Ihara Y, et al. Paternal allelic mutation at the Kcnq1 locus reduces pancreatic β-cell mass by epigenetic modification of Cdkn1c. Proceedings of the National Academy of Sciences. 2015;112: 8332–8337. doi:10.1073/pnas.1422104112

38. Abe-Dohmae S, Ikeda Y, Matsuo M, Hayashi M, Okuhira K, Ueda K, et al. Human ABCA7 supports apolipoprotein-mediated release of cellular cholesterol and phospholipid to generate high density lipoprotein. J Biol Chem. 2004;279: 604–611. doi:10.1074/jbc.M309888200

39. Ikeda F, Deribe YL, Skånland SS, Stieglitz B, Grabbe C, Franz-Wachtel M, et al. SHARPIN forms a linear ubiquitin ligase complex regulating NF-κB activity and apoptosis. Nature. 2011;471: 637–641. doi:10.1038/nature09814

40. Kielar D, Kaminski WE, Liebisch G, Piehler A, Wenzel JJ, Möhle C, et al. Adenosine triphosphate binding cassette (ABC) transporters are expressed and regulated during terminal keratinocyte differentiation: a potential role for ABCA7 in epidermal lipid reorganization. J Invest Dermatol. 2003;121: 465–474. doi:10.1046/j.1523-1747.2003.12404.x

41. Wang N, Lan D, Gerbod-Giannone M, Linsel-Nitschke P, Jehle AW, Chen W, et al. ATP-binding cassette transporter A7 (ABCA7) binds apolipoprotein A-I and mediates cellular phospholipid but not cholesterol efflux. J Biol Chem. 2003;278: 42906–42912. doi:10.1074/jbc.M307831200

42. Satoh K, Abe-Dohmae S, Yokoyama S, St George-Hyslop P, Fraser PE. ATP-binding cassette transporter A7 (ABCA7) loss of function alters Alzheimer amyloid processing. J Biol Chem. 2015;290: 24152–24165. doi:10.1074/jbc.M115.655076

43. Stein CM, Zalwango S, Malone LL, Won S, Mayanja-Kizza H, Mugerwa RD, et al. Genome Scan of M. tuberculosis Infection and Disease in Ugandans. PLOS ONE. 2008;3: e4094. doi:10.1371/journal.pone.0004094

44. Chang K, Deng S, Lu W, Wang F, Jia S, Li F, et al. Association between CD209 - 336A/G and -871A/G Polymorphisms and Susceptibility of Tuberculosis: A Meta-Analysis. PLOS ONE. 2012;7: e41519. doi:10.1371/journal.pone.0041519

45. Naderi M, Hashemi M, Taheri M, Pesarakli H, Eskandari-Nasab E, Bahari G. CD209 promoter –336 A/G (rs4804803) polymorphism is associated with susceptibility to pulmonary tuberculosis in Zahedan, southeast Iran. Journal of Microbiology, Immunology and Infection. 2014;47: 171–175. doi:10.1016/j.jmii.2013.03.013

46. Gay NR, Gloudemans M, Antonio ML, Abell NS, Balliu B, Park Y, et al. Impact of admixture and ancestry on eQTL analysis and GWAS colocalization in GTEx. Genome Biology. 2020;21: 233. doi:10.1186/s13059-020-02113-0

47. Zhong Y, Perera MA, Gamazon ER. On Using Local Ancestry to Characterize the Genetic Architecture of Human Traits: Genetic Regulation of Gene Expression in Multiethnic or Admixed Populations. The American Journal of Human Genetics. 2019;104: 1097–1115. doi:10.1016/j.ajhg.2019.04.009

48. Price AL, Patterson N, Hancks DC, Myers S, Reich D, Cheung VG, et al. Effects of cis and trans Genetic Ancestry on Gene Expression in African Americans. PLOS Genetics. 2008;4: e1000294. doi:10.1371/journal.pgen.1000294

49. H3Africa: current perspectives. 24 Oct 2019 [cited 24 Oct 2019]. Available: https://www.ncbi.nlm.nih.gov/pmc/articles/PMC5903476/

50. Zhao S, Jing W, Samuels DC, Sheng Q, Shyr Y, Guo Y. Strategies for processing and quality control of Illumina genotyping arrays. Brief Bioinform. 2017;19: 765–775. doi:10.1093/bib/bbx012

51. Goldstein JI, Crenshaw A, Carey J, Grant GB, Maguire J, Fromer M, et al. zCall: a rare variant caller for array-based genotyping: genetics and population analysis. Bioinformatics. 2012;28: 2543–2545. doi:10.1093/bioinformatics/bts479

52. Henn Lab SNP-QC pipeline Steps. Henn Lab; 2021. Available: https://github.com/hennlab/snake-SNP_QC

53. Deelen P, Bonder MJ, van der Velde KJ, Westra H-J, Winder E, Hendriksen D, et al. Genotype harmonizer: automatic strand alignment and format conversion for genotype data integration. BMC Res Notes. 2014;7: 901. doi:10.1186/1756-0500-7-901

54. Sudmant PH, Rausch T, Gardner EJ, Handsaker RE, Abyzov A, Huddleston J, et al. An integrated map of structural variation in 2,504 human genomes. Nature. 2015;526: 75–81. doi:10.1038/nature15394

55. Purcell S, Neale B, Todd-Brown K, Thomas L, Ferreira MAR, Bender D, et al. PLINK: a tool set for whole-genome association and population-based linkage analyses. Am J Hum Genet. 2007;81: 559–575. doi:10.1086/519795

56. Delaneau O, Marchini J, Zagury J-F. A linear complexity phasing method for thousands of genomes. Nature Methods. 2012;9: 179–181. doi:10.1038/nmeth.1785

57. Durbin R. Efficient haplotype matching and storage using the positional Burrows– Wheeler transform (PBWT). Bioinformatics. 2014;30: 1266–1272. doi:10.1093/bioinformatics/btu014

58. Kuhn RM, Haussler D, Kent WJ. The UCSC genome browser and associated tools. Brief Bioinform. 2013;14: 144–161. doi:10.1093/bib/bbs038

59. Conomos MP, Reiner AP, Weir BS, Thornton TA. Model-free Estimation of Recent Genetic Relatedness. The American Journal of Human Genetics. 2016;98: 127–148. doi:10.1016/j.ajhg.2015.11.022

60. Alexander DH, Lange K. Enhancements to the ADMIXTURE algorithm for individual ancestry estimation. BMC Bioinformatics. 2011;12: 246. doi:10.1186/1471-2105-12-246

61. Zhou Y, Qiu H, Xu S. Modeling Continuous Admixture Using Admixture-Induced Linkage Disequilibrium. Sci Rep. 2017;7. doi:10.1038/srep43054

62. Uren C, Hoal EG, Möller M. Putting RFMix and ADMIXTURE to the test in a complex admixed population. BMC Genetics. 2020;21: 40. doi:10.1186/s12863-020-00845-3

63. Maples BK, Gravel S, Kenny EE, Bustamante CD. RFMix: a discriminative modeling approach for rapid and robust local-ancestry inference. Am J Hum Genet. 2013;93: 278–288. doi:10.1016/j.ajhg.2013.06.020

64. Dobin A, Davis CA, Schlesinger F, Drenkow J, Zaleski C, Jha S, et al. STAR: ultrafast universal RNA-seq aligner. Bioinformatics. 2013;29: 15–21. doi:10.1093/bioinformatics/bts635

65. Zhong Y, Perera MA, Gamazon ER. On Using Local Ancestry to Characterize the Genetic Architecture of Human Phenotypes: Genetic Regulation of Gene Expression in Multiethnic or Admixed Populations as a Model. bioRxiv. 2018 [cited 18 Feb 2019]. doi:10.1101/483107

66. Stegle O, Parts L, Piipari M, Winn J, Durbin R. Using probabilistic estimation of expression residuals (PEER) to obtain increased power and interpretability of gene expression analyses. Nat Protoc. 2012;7: 500–507. doi:10.1038/nprot.2011.457

67. Raudvere U, Kolberg L, Kuzmin I, Arak T, Adler P, Peterson H, et al. g:Profiler: a web server for functional enrichment analysis and conversions of gene lists (2019 update). Nucleic Acids Research. 2019;47: W191–W198. doi:10.1093/nar/gkz369

